# Delayed transplantation of neural stem cells improves initial graft survival following stroke

**DOI:** 10.1101/2025.02.13.638119

**Authors:** Rebecca Z Weber, Nora H Rentsch, Beatriz Achón Buil, Lina R. Nih, Christian Tackenberg, Ruslan Rust

## Abstract

Neural stem cell therapies hold great promise for improving stroke recovery, but the hostile stroke microenvironment can hinder the initial graft survival. It has long been well documented that the microenvironment evolves over time, making it crucial to identify the optimal transplantation window to maximize therapeutic efficacy. However, it remains uncertain whether acute or delayed local cell transplantations better supports graft viability after stroke. Here, we show that delayed intracerebral transplantation of neural progenitor cells (NPCs) derived from human induced pluripotent cells (iPSCs) at 7 days post stroke significantly enhances graft proliferation and survival, compared to acute transplantation at 1 day post stroke, in a mouse model of large cortical stroke. Using *in vivo* bioluminescence imaging over a 6-week period post-transplantation, we observe a more than 5-fold increase in bioluminescence signal in mice that received delayed NPC therapy, compared to those that underwent acute NPC transplantation. The increased number of cell grafts in mice receiving delayed NPC transplantation was driven by increased proliferation rates early after transplantation, which subsequently declined to similarly low levels in both groups. Notably, we found that the majority of transplanted NPCs differentiated into neurons after 6 weeks, with no significant differences in the neuron-to-glia ratio between acute and delayed transplantation groups. These findings suggest that delayed NPC transplantation improves early graft survival and proliferation, which could help identify the optimal therapeutic window for maximizing the effectiveness of NPC-based therapies in stroke.

## Introduction

Stroke remains a leading cause of disability and death worldwide with no regenerative treatments available.^1^ Stem cell therapy is considered one of the most promising experimental strategies for enhancing stroke recovery.^2^ A promising cell source for cell therapy is neural progenitor cells (NPCs) derived from induced pluripotent stem cells (iPSCs), as they can be expanded indefinitely and differentiated into any cell type, including mature neurons, and glia as shown by us and others previously.^3^ Although local brain transplantation of NPCs has shown preclinical efficacy, the optimal timepoint for transplantation after stroke remains unclear.^4^ While some studies suggest that delayed cell transplantation could be beneficial to avoid the acute inflammatory stroke environment and associated poor cell survival^5,6^, other preclinical and clinical research suggests that cell-based therapies may be most effective when administered as early as 24 h after the stroke onset.^7–9^

We hypothesized that delaying NPC transplantation would enhance survival and tissue integration compared to acute transplantation. To test the hypothesis, we locally transplanted NPCs into mice with large cortical strokes at acute (1 day post-injury, dpi) and delayed (7 dpi) timepoints. Over a 43-day period post-transplantation, we found that NPCs transplanted at 7 dpi displayed higher proliferation and survival rates. Specifically, the proliferation rate was notably higher in the early days following transplantation, suggesting that the hostile stroke environment limits the acute growth of NPCs. Furthermore, we observed that most transplanted cell grafts successfully differentiated into mature neurons and glia within six weeks. The neuron-to-glia ratio remained consistent across both acute and delayed transplantation groups, indicating no significant effect of transplantation timing on NPC fate commitment.

## Results

To determine the survival and proliferation of the cell grafts, we locally transplanted NPC expressing firefly luciferase (rFluc) in immunodeficient Rag2^−/−^ mice at 1 day (NPC_acute_) or 7 days (NPC_delayed_) post-stroke (**Fig.1A**).

We first confirmed in all mice that the stroke induction was successful and consistent across all mice, resulting in at least 50% reduction in cerebral blood flow two hours after induction compared to the contralesional site (p < 0.0001) (**Fig. 1B, C**). Mice were then randomly allocated to the experimental groups with a similar stroke severity across groups. Next, we analyzed longterm survival of rFluc-expressing NPC_acute_ and NPC_delayed_ throughout the time course of 6 weeks after stroke via bioluminescence imaging (**Fig. 1D**). After 43 days post stroke, we observed a considerably stronger bioluminescence signal (NPC_acute_: 3.2 ph/s/cm3/sr x 10^5^) vs NPC _delayed_: 60 ph/s/cm3/sr x 10^5^, p < 0.05) (**Fig. 1E**). Additionally, a notable decrease in bioluminescence signal was observed within the first 3 days after transplantation, with a reduction of 56% in the NPC_acute_ group compared to 51% in the NPC_delayed_ group (**Fig. 1F**).

**Figure 1.**
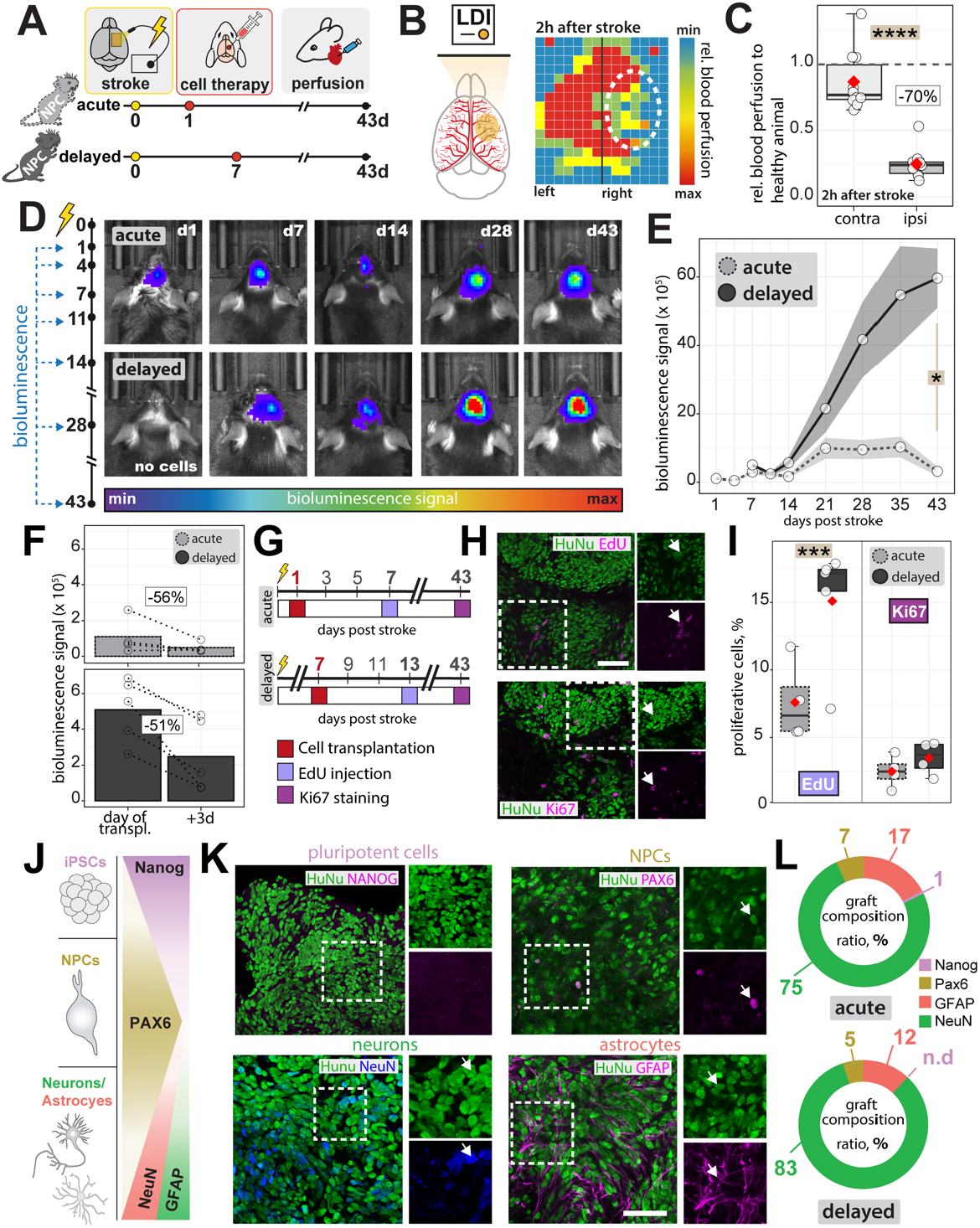
Delayed transplantation of NPCs improves long-term graft survival post-stroke. (A) Schematic illustration showing the experimental design. Immunodeficient Rag2^−/−^ mice underwent stroke induction followed by local transplantation of rFluc-expressing NPCs at either 1 dpi (acute) or 7 dpi (delayed). (B) Laser Doppler imaging confirms reduced cerebral blood flow (CBF) after stroke. (C) Quantification of CBF 2 hours after stroke induction. (D) Representative bioluminescence imaging (BLI) illustrating NPC survival over 6 weeks in both groups of selected time points. (E) Quantification of BLI signal over time and (F) within the first 3 days after transplantation in both groups. (G) Schematic timeline of proliferation assessment using EdU incorporation at 7 days post-transplantation and Ki67 staining at 42 days (acute) and 35 days (delayed) post-transplantation to track graft proliferation. (H) Representative immunofluorescence images (scale bars: 50 µm) and (I) quantification of EdU^+^ NPCs at seven days post-transplantation and Ki67^+^ NPCs at 35 dpi (delayed) and 42 dpi (acute) days in both groups. (J) Illustration showing phenotyping panel with pluripotency marker Nanog, NPC marker PAX6, neuronal marker NeuN, and astrocytic marker GFAP. (K) Representative immunofluorescence images of grafted NPCs (HuNu+) at six weeks post-transplantation. Scale bars: 50µm. (L) Quantification of graft composition in acute vs. delayed transplantation groups. Data are shown as mean distributions where the red dot represents the mean. A total of N = 8 animals were used, with 4 animals per group. Boxplots indicate the 25% to 75% quartiles of the data. For boxplots: each dot in the plot represents one animal. Line graphs are plotted as mean ± SEM. Significance of mean differences was assessed using an unpaired Mann-Whitney U Test (C and E) or an unpaired t-test (I). Statistical significance was set at *, p < 0.05; **, p < 0.01; ***, p < 0.001.

These findings suggest that delayed NPC transplantation results in a higher detectable graft signal over six weeks, indicating potential benefits for graft survival and proliferation after stroke.

Next, we aimed to determine whether the observed differences in bioluminescence signal were due to increased proliferation early after transplantation or at later stages. To investigate this, we injected the nucleotide analog EdU to label proliferating cells at 7 days post-transplantation and conducted Ki67 staining after tissue collection (**Fig. 1G-H**). We detected a twofold increase in EdU proliferation within the graft in NPC_delayed_ compared to NPC_acute_ one week post transplantation (15.1% vs 7.6%; p < 0.001, **Fig 1I**). Notably, NPC proliferation levels were at similar low rates at 43 days after stroke (NPC_acute_: 2.5% Ki67^+^ cells vs NPC_delayed_: 3.5% Ki67^+^ cells, p > 0.05, **Fig 1I**).

Most transplanted NPCs differentiate into astrocytes and neurons within the first month, as shown in previous studies.^10^ To identify whether the transplantation timing after stroke can affect the graft composition, we phenotyped human nuclei (HuNu^+^) positive cell grafts for pluripotent stem cell (Nanog), NPC (Pax6), mature neuronal (NeuN) and astrocytic (GFAP) markers. We observed that the majority of cell grafts differentiate into NeuN^+^ mature neurons (NPC_acute_: 75% vs NPC_delayed_: 83%), with only a small fraction of grafts expressing astrocytic GFAP^+^ (NPC_acute_: 17% vs NPC_delayed_: 12%), and Pax6 (NPC_acute_: 7% vs NPC_delayed_: 5%).

These findings suggest that delayed NPC transplantation promotes an initial surge in proliferation rates, possibly through a less hostile stroke environment. The timing of transplantation does not seem to affect overall differentiation patterns, as both acute and delayed grafts consistently maintain a similar neuronal-to-astrocytic ratio, with no detectable pluripotent residuals.

## Discussion

In this study, we show that delayed (7 dpi) transplantation of NPCs significantly improves graft proliferation and survival compared to acute (1 dpi) transplantation in a photothrombotic stroke model.

In the acute phase following stroke, the cerebral microenvironment is characterized by highly inflammatory responses, oxidative stress and a disrupted blood brain barrier (BBB), thus creating a hostile setting for cell therapy. As early as 24 hours post-stroke, reactive oxygen species (ROS) and a plethora of inflammatory mediators (TNF-α, IL-1β, IL-6, etc.) compromise the BBB, leading to infiltration of immune cells (neutrophils, macrophages and T cells) into the brain parenchyma.^11^ At the same time, matrix metalloproteinases (MMPs) degrade perivascular basement membranes and tight junction proteins, further increasing BBB permeability, enabling the entrance of toxic substances.^12,13^ These factors not only induce direct cellular damage but also might hinder the integration and differentiation of newly transplanted cells. After the early activation of the immune system, the inflammatory milieu partially subsides^14^, the BBB regains integrity^15,16^ and oxidative stress levels diminish^17^. Thus, the subacute phase may offer a more favorable environment for the survival and engraftment of transplanted NPCs, compared to the highly hostile conditions that prevail during the acute phase.^4^ In support of these assumptions, previous studies have reported that delayed transplantation following lesion ameliorated graft survival rates. Embryonic motor cortical neuron grafts showed significantly enhanced vascularization, survival, and transplant-to-host projections when transplanted 1 week after injury, compared to earlier transplantation timepoints.^18^ And delivery of bone marrow-derived MSCs between 2- and 7-days post-stroke significantly increased outcome compared to administration 24h after injury.^19^

While some previous work indicates that acute systemic transplantation of stem cells might be beneficial during acute time window^20,20^, these effects likely are observed through paracrine signaling within the peri-infarct region (e.g., reduced inflammation, improved angiogenesis)^21–23^ or peripheral interactions that modulate the repair process primarily outside the brain.^24–26^

Future studies should explore the mechanisms underlying the enhanced survival observed in grafts transplanted at delayed timepoints (7dpi), particularly by investigating how inflammatory modulation and niche adaptation contribute to host–graft integration. Additionally, cell priming strategies, such as exposing NPCs to hypoxia or hypothermia, pre-conditioning cells with growth factors, gene-editing to promote the expression of protective genes, modulating the immune response, or providing a supportive scaffold, should be explored to determine whether these approaches improve graft survival, engraftment, and proliferation within the subacute stroke environment. Indeed, these strategies have been shown to trigger adaptive responses in cells, such as increased growth factor secretion and improved stress tolerance, which may enhance NPC viability and integration in the injured brain. Understanding the molecular mechanisms behind these approaches could help optimize cell-based therapies for better functional recovery after stroke.

Interestingly, we did not observe differences in the graft composition. While few previous studies have suggested that hypoxic changes can alter the glia vs neuron fate commitment in NPC to the glial phenotype^27–29^, we did not observe a significant change in the graft composition. Potentially, these findings may indicate that the stroke microenvironment do not exhibit strong differences in O_2_ availability at 1 and 7 days post injury, but instead reflects a reduced inflammatory state, which has been observed before in stroke patients.^30^

While our results show that the overall number of NPCs increases over six weeks when cell grafts are transplanted one week after injury, it is important to note that this enhanced bioluminescent signal does not indicate tumor formation. In a previous study, we found no histological evidence of tumorigenesis and instead observed that most grafted NPCs differentiated into neurons, as evidenced by the high positivity for the neuronal markers NeuN, MAP2, and βIII-tubulin.^3^ Future work could look at more chronic time points after stroke and address whether the increased number of cells will impact long-term functional recovery. It will also be important to characterize if and how the timing of transplantation impacts the degree of graft vascularization and the extent of axonal projections formed by the transplanted neurons, as both parameters are crucial for establishing long-term functional recovery.

From a clinical standpoint, delayed transplantation offers several advantages, including improved patient stabilization, a potentially better prediction of stroke injury progression, and the opportunity to extend the currently limited 4–24-hour therapeutic window.

Overall, our findings suggest that delaying NPC transplantation by one week may enhance the therapeutic potential of NPC-based therapies in stroke and potentially other related neurological disorders.

## Methods

### Animals

All animal experiments were conducted at the Laboratory Animal Services Center (LASC) in Schlieren in compliance with local regulations for animal research and approved by the Veterinary Office in Zurich, Switzerland. A total of eight (four male and four female) adult genetically immunodeficient Rag2^-/-^ mice (20–30 g) were used for this study. The group sizes were determined by power calculations and prior studies.^3,10,31,32^ Mice were housed at the LASC in top-filter laboratory cages under OHB conditions in a temperature- and humidity-controlled environment with a 12/12-hour light/dark cycle (lights on from 6:00 a.m. to 6:00 p.m.). Animals were housed in groups of two to four per cage and had ad libitum access to standard diet food pellets and water.

### Cell Culture

For our cell-based stroke therapy, we have developed a chemically defined, xeno-free differentiation protocol to generate neural progenitor cells (NPCs) from human iPSCs. Gene and protein expression analysis confirmed the stable upregulation of NPC markers (Pax6, Sox1, Nestin) and the downregulation of pluripotency markers (Oct4) over 15 passages, indicating successful differentiation. Before transplantation, NPCs were transduced with a lentiviral vector encoding a bioluminescent (red firefly luciferase) and fluorescent (eGFP) reporter, as previously described^3^, enabling *in vivo* tracking of transplanted cells.

### Photothrombotic stroke induction

Photothrombotic stroke was induced as previously described ^15,33–35^. Anesthesia was initiated with 4% isoflurane in oxygen and maintained at 1–2% using a self-fabricated face mask. Mice were positioned in a stereotactic frame (David Kopf Instruments), and body temperature was maintained at 36–37°C using a heating pad. Anesthesia depth was confirmed by the absence of a toe-pinch reflex, and eye lubricant (Vitamin A, Bausch & Lomb) was applied to prevent corneal drying. The head was shaved, and Emla™ Creme 5% (lidocaine/prilocaine) was applied to the scalp and ears before inserting the ear bars for skull fixation. A ∼1 cm midline incision was made to expose bregma and lambda, and the periosteum of the lower right hemisphere was scraped off. The exact stroke induction site [-2.5mm to +2.5mm medial/lateral and 0mm to +3mm from bregma] was identified under an Olympus SZ61 surgical microscope and marked. Rose Bengal (15 mg/ml in 0.9% NaCl) was injected intraperitoneally (10 μl/g body weight) 5 minutes before illumination to allow systemic circulation. The marked cortical region was illuminated for 10 minutes using an Olympus KL1500LCD cold light source (150W, 3000K, 4mm × 3mm surface area). After illumination, mice were placed in a recovery cage, and stroke induction was confirmed 30 minutes post-surgery using laser Doppler imaging (LDI). The incision was sutured using a Braun surgical suture kit, disinfected with Betadine®, and animals were monitored under standard post-operative care.

### NPC transplantation

Transplantation of neural progenitor cells (NPCs) was performed as previously described. The surgical procedure followed the photothrombotic stroke model up to the point of illumination. Animals were divided into three groups: NPC transplantation 1 day post-stroke (acute), NPC transplantation 7 days post-stroke (delayed), and a vehicle control group.

After identifying bregma, the stereotactic device was adjusted to the injection site coordinates [AP: +0.5 mm, ML: +1.5 mm, DV: −0.6 mm relative to bregma]. A small ∼0.8 mm hole was drilled into the skull, ensuring penetration through the cortical bone. A 10 μl Hamilton syringe fitted with a 30G needle containing 2.5 μl of NPC suspension (1×10^5^ cells/μl) was mounted onto the stereotactic device and positioned above the injection site. The needle was slowly lowered to the calculated depth, with an additional 0.05 mm penetration to create a small cavity, preventing cell suspension spillover.

Once retracted to the target depth, NPCs were injected at a steady rate of 2 nl/s into the brain parenchyma. After injection, the needle remained in place for 5 minutes to allow cell settlement before being withdrawn slowly. The cavity was sealed with Histoacryl®, and once hardened, the skin was sutured. Animals were returned to an empty cage and monitored under standard post-operative care.

### Bioluminescent imaging (BLI)

Animals were imaged using the IVIS® Lumina III In Vivo Imaging System, with the head region carefully shaved using an electric razor to ensure optimal signal detection. Mice received an intraperitoneal injection of D-luciferin (300 mg/kg body weight) dissolved in PBS, which was sterile-filtered through a 0.22 μm syringe filter. Bioluminescence images were acquired at multiple time points (1, 3, 7, 14, 21, 28, 35 and 43, days post-injury) to monitor transplanted cell survival and distribution.

Imaging data were analyzed using Living Image software (V 4.7.3). A region of interest (ROI) (height, width = 1.8 cm) was positioned directly over the color-coded signal between the ears and nose of each animal. Two additional ROIs were placed: one at the posterior body region to quantify background signal and another in a randomly selected area proximal to the body to measure image noise. Bioluminescence intensity was quantified as total photon flux (photons per second). Data was recorded in Microsoft Excel and visualized using R software for statistical analysis.

### Tissue processing and immunohistochemistry

Animals were transcardially perfused with Ringer solution (0.9% NaCl). Brain tissue was then extracted and immediately frozen for immunohistochemistry. Animals were perfused first with Ringer solution, followed by 4% PFA, and brains were subsequently incubated in 4% PFA for 6 hours. Coronal brain sections (40 μm) were prepared using a Thermo Scientific HM 450 microtome, washed in 0.1M phosphate-buffered saline (PBS), and blocked for 1 hour at room temperature in 500 μl of blocking buffer containing 5% donkey serum diluted in 1× PBS + 0.1% Triton X-100. Sections were then incubated overnight at 4°C on a Unimax 1010 shaker (∼90 rpm) with the primary antibody Ki-67 (rabbit, 1:200, Invitrogen) to assess cell proliferation. The following day, sections were washed in PBS and incubated with the appropriate secondary antibody for 2 hours at room temperature, followed by staining with DAPI (Sigma, 1:2000 in 0.1M PBS). Finally, sections were mounted onto Superfrost Plus™ microscope slides and coverslipped using Mowiol® mounting solution.

To further evaluate cell proliferation 1 week after cell transplantation, 5-ethynyl-2′-deoxyuridine (EdU) labeling was performed. EdU was dissolved in sterile PBS at a concentration of 5 mg/ml and administered via intraperitoneal injection at 50 mg/kg body weight 7 days after transplantation. Brain sections were incubated for 30 minutes with a reaction mix buffer (1× per well: 325 μl ddH_2_O, 25 μl 2M Tris, 100 μl 10mM CuSO_4_, 0.1 μl 10mM AlexaFluor 647, 100 μl 500mM ascorbic acid). Sections were then washed with 0.1M PBS, counterstained with DAPI, mounted onto Superfrost Plus™ microscope slides, and coverslipped with Mowiol® solution.

### Statistical analysis

Statistical analysis was performed using RStudio (Version 4.04). Sample sizes were designed with adequate power in line with previous studies from our group and relevant literature. Data normality was assessed using the Shapiro-Wilk test. Post hoc multiple comparisons were corrected using the Bonferroni method. For bioluminescence imaging analysis, signal intensity was compared day-wise using an unpaired Mann–Whitney *U* test. Proliferative cell counts assessed via EdU and Ki67 staining were compared between acute and delayed transplantation groups using an unpaired two-sided t-test. Blood flow measurements obtained by Laser doppler were compared between healthy and injured hemisphere using an unpaired Mann– Whitney *U* test. Box plots display median, interquartile range (IQR), and individual data points. Line graphs are expressed as mean ±SEM. Statistical significance was set at p < 0.05.

### AI assistance disclosure

Grammar and typos were corrected with the assistance of ChatGPT V 4o (OpenAI). No content generation or data analysis was conducted using AI.

## Data availability

All data is available upon request.

## Ethics declaration

All in vivo experiments were performed at the Laboratory Animal Services Center (LASC) in Schlieren, Switzerland according to the local guidelines for animal experiments and were approved by the Veterinary Office of the Canton Zurich in Switzerland (Protocol number ZH209/2019 and ZH110/2023).

## Acknowledgements

-

## Author contribution

RZW, CT, RR contributed to overall project design. RZW, NHR, LN, BAB, CT, RR conducted and analyzed experiments. RZW, RR made figures. CT, RR supervised the study. RZW, LN, CT, RR wrote and edited the manuscript with input from all authors. All authors read and approved the final manuscript.

